# RPnet: A Reverse Projection Based Neural Network for Coarse-graining Metastable Conformational States for Protein Dynamics

**DOI:** 10.1101/2021.08.04.455071

**Authors:** Hanlin Gu, Wei Wang, Siqin Cao, Ilona Christy Unarta, Yuan Yao, Fu Kit Sheong, Xuhui Huang

## Abstract

Markov State Model (MSM) is a powerful tool for modeling the long timescale dynamics based on numerous short molecular dynamics (MD) simulation trajectories, which makes it a useful tool for elucidating the conformational changes of biological macromolecules. By partitioning the phase space into discretized states and estimate the probabilities of inter-state transitions based on short MD trajectories, one can construct a kinetic network model that could be used to extrapolate long time kinetics if the Markovian condition is met. However, meeting the Markovian condition often requires hundreds or even thousands of states (microstates), which greatly hinders the comprehension of conformational dynamics of complex biomolecules. Kinetic lumping algorithms can coarse grain numerous microstates into a handful of metastable states (macrostates), which would greatly facilitate the elucidation of biological mechanisms. In this work, we have developed a reverse projection based neural network (RPnet) method to lump microstates into macrostates, by making use of a physics-based loss function based on the projection operator framework of conformational dynamics. By recognizing that microstate and macrostate transition modes can be related through a projection process, we have developed a reverse projection scheme to directly compare the microstate and macrostate dynamics. Based on this reverse projection scheme, we designed a loss function that allows effectively assess the quality of a given kinetic lumping. We then make use of a neural network to efficiently minimize this loss function to obtain an optimized set of macrostates. We have demonstrated the power of our RPnet in analyzing the dynamics of a numerical 2D potential, alanine dipeptide, and the clamp opening of an RNA polymerase. In all these systems, we have illustrated that our method could yield comparable or better results than competing methods in terms of state partitioning and reproduction of slow dynamics. We expect that our RPnet holds promise in analyzing conformational dynamics of biological macromolecules.

## I. INTRODUCTION

Many biologically relevant events occur at milliseconds or longer timescales.^1–3^ While molecular dynamics (MD) simulation tools and computation resources had a large advancement in the past few decades, obtaining a reliable sampling of these long-timescale events is still challenging. Markov State Model (MSM) is a mathematical framework that was built from a large number of short MD simulations but allows the estimation of long timescale dynamics, so that conformational sampling could be done in an highly parallelized manner.^4–27^ MSM works by decomposing the phase space into discrete regions (“states”) and then estimating the interstate transition probabilities at a specified lag time. If the complex system dynamics has a separation of timescale and the resulting transition probability matrix (TPM) satisfies the Markovian (memoryless) condition, the population evolution of the system under study could be calculated through repeated self-propagation of the TPM. In order to construct a Markovian model with an affordable lag time that allows efficient sampling (bound by the length of MD simulations to estimate transition probabilities), hundreds or thousands of microstates are often needed, but the large number of microstates present would hinder the interpretation of biological insights. This is a dilemma often encountered in MSM construction that models with large number of states are often hard to interpret, while models with only a few states are much more challenging to meet the Markovian condition.

One popular way to strike the balance and obtain an MSM with only a handful of states is via a two-stage procedure.^8,28,29^ Collective variables that can properly describe the conformational dynamics of biological macromolecules are first chosen (e.g. by the time-lagged independent component analysis: tICA^5,30^), all the conformations are then split into a large number of “microstates” using geometric clustering algorithms like k-centers^7,31^ or k-means^32,33^ clustering based on the chosen representation, giving rise to a microstate MSM. It is then followed by a kinetic lumping procedure, merging the microstates into a handful coarse-grained metastable states (macrostates), so that we can better interpret the biological mechanisms.^2,3,34^ One widely used group of kinetic lumping algorithms makes use of the dominant eigenvectors of the TPM to determine the lumping, like the Perron-Cluster Cluster Analysis (PCCA)^35^ that repeatedly bi-partitions the microstates into groups based on the sign structures of the dominant eigenvectors, or its robust variants.^36–38^ There are also methods based on Bayesian inferences, as exemplified by the Bayesian agglomerative clustering engine (BACE),^39^ by repeatedly merging the microstate pairs with smallest BACE Bayes factor. In another Bayesian kinetic lumping algorithm, a Gibbs sampling algorithm is applied to facilitate the search of the optimal kinetic lumping.^40^ Methods that explicitly considers the possible transition paths in the microstate transition matrix also exists, an example of which is the Most Probable Path (MPP) lumping algorithm.^41^

Rather than the two-step procedure discussed above, several deep-learning methods have recently been developed to obtain macrostate models directly from MD simulations trajectories.

A notable example is the VAMPnet,^42^ which takes Cartesian coordinates or other physical variables extracted from MD simulation trajectories as input, and directly produces a macrostate assignment and the corresponding Markov model through a neural network. The quality of lumping is determined by the VAMP-2 score based on the variational principle of the conformational dynamics. There is also a related approach with a symmetrization constraint to enforce the detailed balance, known as State-free Reversible VAMPnet (SRV),^43^ which enforces the detailed balance condition, which has been successfully applied to the study of Trp-cage protein.^44^

The kinetic lumping can also be viewed as a projection process of conformational dynamics. This view makes use of the projection operator framework in statistical mechanics developed by Zwanzig and Mori^45–47^. In fact, this idea has been previously realized by Hummer and Szabo in multistate kinetics, where the original dynamics of the system is projected into a reduced system by keeping the exact dynamics of the reduced system in both non-Markovian and Markovian region.^48^ We have also applied such a projection scheme to understand the conformational dynamics of complex systems through the extrapolation of occupancy-number correlation based on a given macrostate partitioning.^29^ A similar idea with the projection operator scheme is to examine the transfer operator^6^ itself as opposed to population correlation, which also provides a useful framework for developing the kinetic lumping algorithm.

Based on the above view of the projection scheme for conformational dynamics^29^, we have developed a novel deep learning-based kinetic lumping method: a reverse projection based neural network (RPnet) method. The key insight of RPnet lies in its loss function that can quantify the quality of lumping via a reverse projection process of conformational dynamics. The reverse projection scheme utilizes the projection operator to evaluate macrostate models, by assessing the ability to match the dynamics of the microstate model in the macrostate resolution. In this scheme, an overlapping matrix between microstate eigenvectors (i.e. the eigenvectors of the microstate TPM) and a backward projected vector of macrostate eigenvectors is adopted as the scoring function for evaluation. In the RPnet, this eigenvector-based scoring function is used as the loss function, and the optimization of this loss function would improve the matching of dynamics between macrostate models and the microstate model. This design of the loss function makes RPnet different from other deep learning-based lumping methods such as VAMPnet, where the loss function is designed based on the variational principle of conformational dynamics. In our loss functions, we have formulated a reverse projection scheme to allow direct comparisons between the dynamics of lumped macrostate models and the original microstate models to evaluate the quality of lumping, and further optimize the macrostate boundaries. In terms of architectures of deep learning networks, we have constructed a two-lobe encoder neural network. We demonstrate that our RPnet method performs well when applied to study systems ranging from numerical potentials to the complex RNA polymerase (RNAP) system. We anticipate that our RPnet holds promise to be widely applied to study biomolecular dynamics.

## II. METHODS

### A. Kinetic lumping as a projection operation

To understand the basis of our design, we first examine the relationship between populations of microstates (small states, usually in a number of hundreds to thousands) and macrostates (large metastable states, often only a handful of macrostates), and examine how the populations evolve under the projection operator framework.

The transition probability matrix (TPM) of the microstate model (**T**(*τ*)) and the macrostate model (**M**(*τ*)) for a prespecified lagtime *τ* are defined as follows,

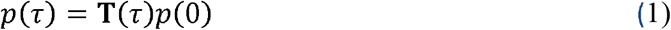

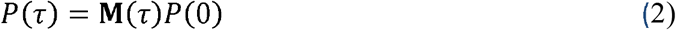

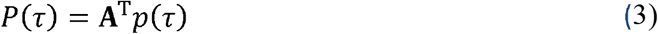

where *p* and *P* are vectors (with dimensions *n* and *N* respectively) corresponding to the microstate and macrostate populations, respectively. **A** is the matrix defining the mapping from the microstates (*j*) to macrostates (*I*): **A**_*jI*_ = 0 if j ∉ I or **A**_*jI*_ = 1 if j ∈ I, thus **A** reflects the macrostate lumping of the system. We note that left-multiplication of **A**^T^ to the microstate population vector is effectively summing up all the populations of microstates within each of the macrostates. Combining Eq (1)-(3), we show that the microstate and macrostate transition probability matrices can be related by the following equations:

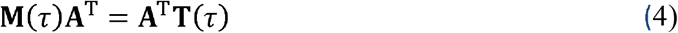

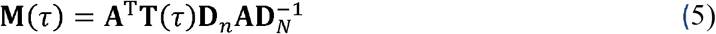

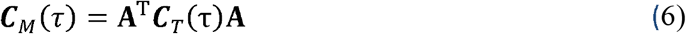

where *C*_*M*_ and *C*_*T*_ correspond to transition count matrices of macrostates and microstates respectively, and **D**_*n*_ and **D**_*N*_ are diagonal matrices containing the microstate and macrostate equilibrium populations. Eq. (5) is straightforwardly obtained as 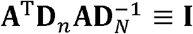 is always an identity matrix, following the fact that **D**_*n*_ and **D**_*N*_ is related by summing up all the equilibrium populations within the corresponding macrostates. We will then examine the “evolution of population” at microstate and macrostate level (*p*(*τ*) and *P*(*τ*), respectively). Under the Markovian condition, with the lag time of *τ*, propagating the microstate population for *m* times using the transition matrix **T**(*τ*) should give the population at time *mτ*:

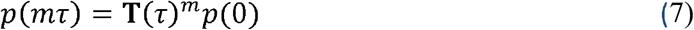

The minimum lag time *τ* for which the population is well described by the equation above is called the Markovian lagtime time *τ*_0_ of the kinetic network model.

The coarse-grained macrostate model is obtained by grouping a number of microstates into one macrostate, where the population of a coarse-grained macrostate is the sum of all corresponding microstates,

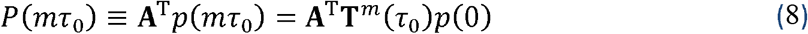

This equation also implies that the population distribution *P* of coarse-grained macrostate at time *t* = *mτ*_0_ could be obtained through repeated propagation of the original (microstate) population by **T**(τ_0_).

One important point to note is that, although the evolution of *p*(*t*) is Markovian, the evolution of *P*(*t*) is not necessarily so, because information of all elements in *p*(0) (as opposed to only information of *P*(0)) is needed in order to obtain *P*(*t*), as seen in Eq (8). Specifically, even if the macrostate population distribution *P*(0) is known, the given population vector could actually be mapped to many different microstate distributions *p*(0), because the population distribution within each macrostate could be assigned in different ways. Although these distributions could all be mapped to the same *P*(0), the intra-macrostate relaxation processes are different, and this would subsequently affect the evolution of macrostate dynamics. Therefore, obtaining a Markovian propagation of *P*(t) is more challenging.

We note that the “lumping” process can be viewed as a projection, by making use of the projection operator framework that was previously developed by Szabo and Hummer.^48^

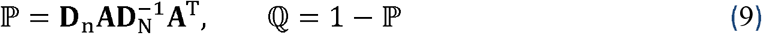

where **D**_*n*_ is the diagonal matrix containing the population of the original kinetic network model and **D**_*N*_ denotes the coarse-grained one. Because the coarse-grained population is obtained by summing the populations of the states in the original model, **D**_*N*_ = **A**^T^ **D**_*n*_ **A**, we can then easily prove the idempotency ℙ^2^= ℙ, ℚ^2^ = ℚ, ℙℚ = 0, and also the identity **A**^T^ ℙ≡**A**^T^. (See Appendix A for the proof).

The projection operator then relates the population *P*(*t*) at the macrostate level and population *p*(*t*) at the microstate level.

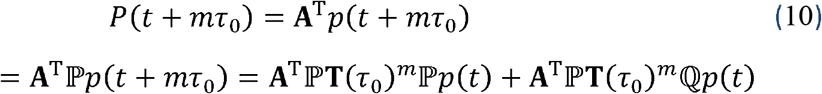

### B. Evaluating the Quality of Kinetic lumping via reverse projection

To closely connect the dynamics of microstate and macrostate models, we have formulated a “reverse projection” process by mapping eigenvectors of the macrostate TPM (N transition modes) back to the microstate space (n eigenvectors, *n* ≫ *N*). The “reverse projected” transition mode of a given lumping matrix **A** is defined as.

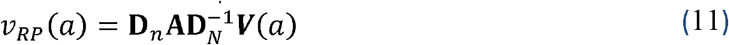

where 1 ≤ *a* ≤ *N*, meaning that only the top *N* modes are considered. Properties of this reverse projection can be found in Appendix A.

To better illustrate the actual meaning of the “modes” other than its mathematical representations, we have made an illustration using a simple 1D potential (Fig. 1). When there is a good lumping where all four energy minima in the 1D potential are properly identified as metastable states (Left panel of Fig. 1b), the “reverse projected” modes 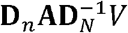 (Green curves in right panels of Fig. 1b) agree well with the original transition modes *v* (Fig. 1a). In sharp contrast, when there is a bad lumping (Left panel of Fig. 1c), the reverse-projected modes (Red curves in right panels of Fig. 1c) largely deviate from the original transition modes. In addition, when we examine closely the reverse projected modes (green lines in Fig. 1b and red lines in Fig. 1c), we notice the reverse-projected modes are “continuous” within each macrostate region, where there are clear discontinuities at the boundaries of macrostates (especially when a “bad lumping” was evaluated in Fig. 1c). This could be understood from the definition of the reverse projection (Eq (*11*)), that the components of the reverse projected modes within each macrostate are proportional to the equilibrium population within the macrostate. This simple 1D potential example clearly illustrates that the resemblance between microstate transition modes and reverse projected modes could serve as a way to examine the quality of state boundaries.

**FIG. 1.**
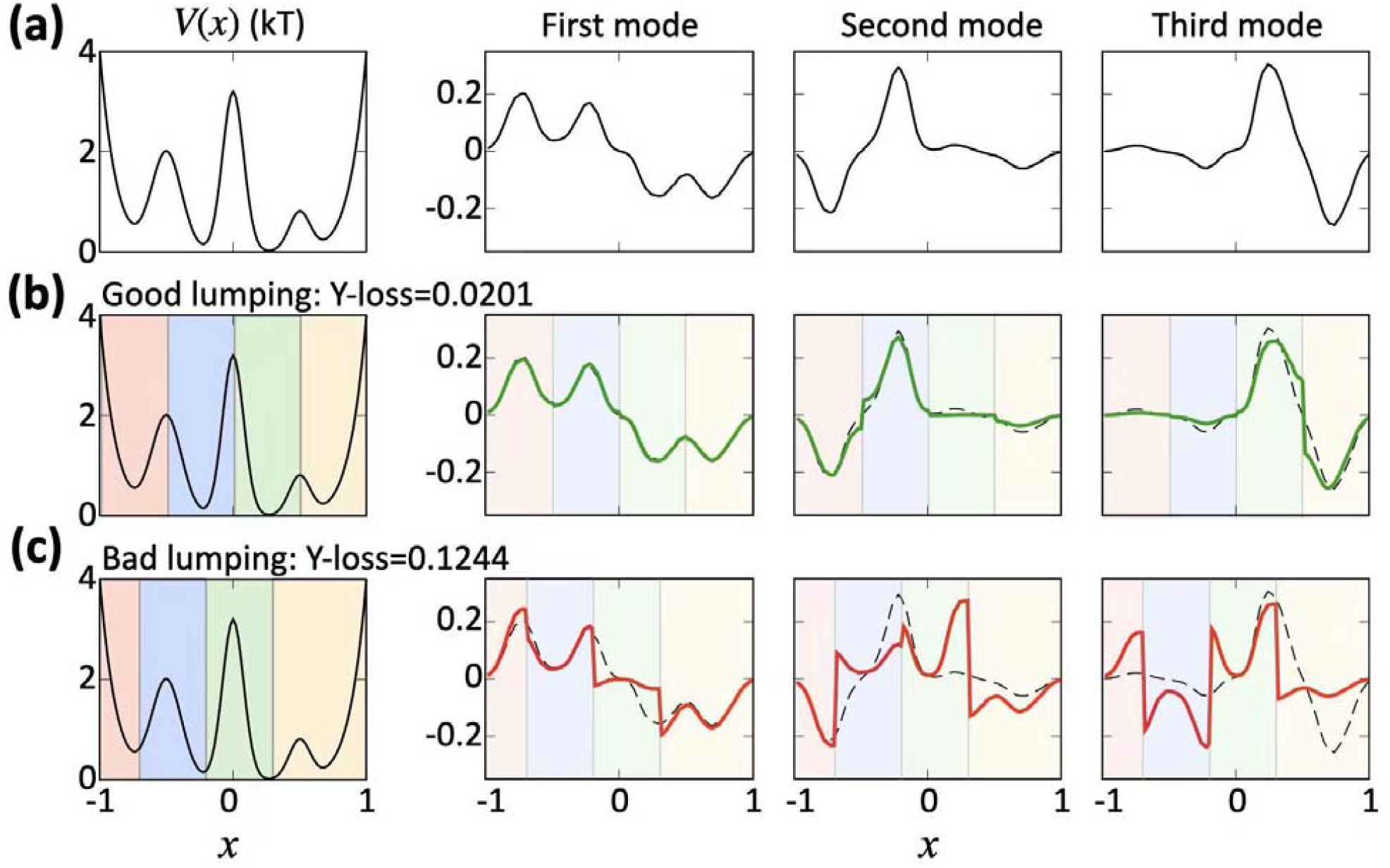
Illustration of reverse projected modes in good and bad lumping. (a) Energy landscape *V*(*x*) and the corresponding microstate transition modes. (b) Reverse projected modes of good lumping. (c) Reverse projected modes of bad lumping. The dashed line in (b) and (c) are the same as the microstate transition modes in (a), shown for the ease of comparison. It is clear from the figure that the reverse projected modes are smooth within each macrostate region, but at the boundaries between two macrostates, clear discontinuities could be present, especially for the modes corresponding to a bad lumping.

To assess the quality of lumping, we quantify the similarity between the transition modes of the microstate models and “reverse projected” modes. Thus, we have defined the following overlap matrix **Y** to quantify the similarity,

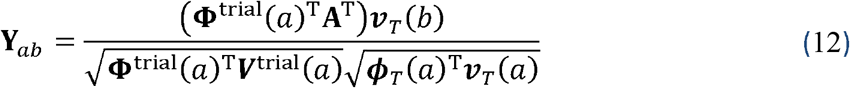

Where ***V***^trial^ and **Φ**^trial^ denotes the right and left eigenvectors of the TPM of the proposed macrostate lumping, respectively, and ***v***_*T*_ and ***ϕ***_*T*_ denotes the right and left eigenvectors of the original microstate transition matrix **T**. Basically, **Φ**^trial^(*a*)^T^**A**^T^ is essentially the left reverse-projected eigenvector for the trial macrostate partitioning (refer to Appendix A for details), and the denominator is the normalization factor. **Y**_*ab*_ therefore reflects the overlap between the reverse projected transition modes and the original microstate transition modes.

Because matrix **Y** measures overlap of the original and the reverse projected eigenvectors, and it is clear from Fig. 1 that the resemblance between the two would be useful for directly quantifying the quality of state boundaries. Matrix **Y** is therefore a good candidate as a loss function for automatic optimization of state boundaries. This loss function will be different from the popular methods such as VAMP-2 score that are based on variational principle of the conformational dynamics. Basically, a perfectly Markovian lumping will be seen with the matrix **Y** equals to identity matrix (see Appendix A for details), so we may take the Freboneus norm of the difference between the **Y** matrix against an identity matrix

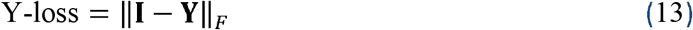

to serve as a loss function for optimization purpose (referred to as Y-loss hereafter). This proposal is also consistent with the lumpability condition proposed by Kemeny and Snell,^49^ who have introduced the “lumpability” condition to reduce the size of state space of some continuous-time Markov chains in probability theory. We can see in Fig. 1 that Y-loss can indeed clearly distinguishes the good from bad lumping, as the good lumping in Fig. 1b has a much smaller Y-loss than the bad lumping in Fig. 1c.

### C. Optimizing Kinetic lumping using an encoder neural network

Note that the minimization of ‖**I** − **Y**‖_*F*_ with respect to the choice of lumping **A** is a highly non-linear problem. We have thus designed a neural network for this optimization. The “lumping” of microstates into macrostates is understood as a “membership assignment”, in which the input of microstate assignment (as a 1 × *n* one-hot vector, where n is the number of microstate) is mapped to the macrostate assignment (as a 1 × *N* membership vector, where *N* is the number of microstate), and we aim to search for a (fuzzy) state assignment by finding a good membership assignment seen as an encoder network. At the end, a crisp lumping is obtained by taking the assignment with maximum membership.

Our algorithm basically consists of four steps (see Fig. 2(a)):

**FIG. 2.**
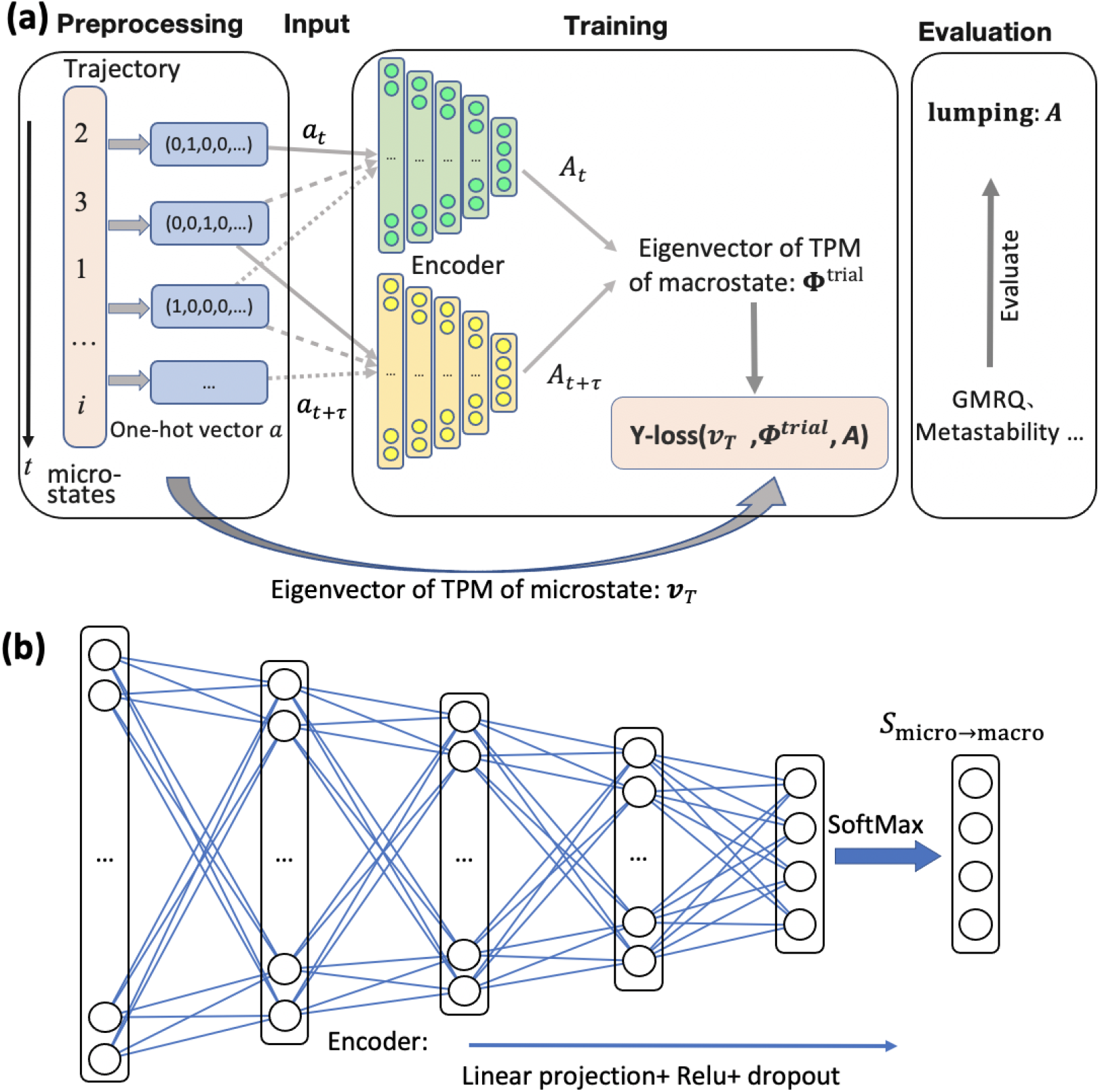
Architecture of the RPnet. (a) Overall architecture of the RPnet. (b) Details of the encoder network.

1. **Preprocessing**: Computing the microstate Transition Count Matrix (TCM) by counting the number of transitions between prespecified states. And converting every frame of microstate assignment (from one or many trajectories) into 1 × *n* one-hot vectors.
2. **Input**: Feeding pairs of one-hot vector assignments (separated by a prespecified lagtime) into two encoder networks that share same architecture and weights. The output of each network is a 1 × *N* vector, representing the membership of the input microstate to the *N* macrostates. The output is also used to compute the macrostate TCM.
3. **Training**: Based on the microstate and macrostate TCM, computing the Y-loss (Eq. (13)), and using backpropagation to optimize the network until convergence.
4. **Evaluation**: The optimized lumping matrix is computed by enumerating all the microstate one-hot vectors and stacking the corresponding output (resulting in an matrix). The quality of the output is also examined by other criteria, including but not limited to metastability or generalized matrix Rayleigh quotient (GMRQ).^50^

More details of the algorithm can be seen from the pseudocode in Appendix B.

### D. Architecture of the encoder network of RPnet

The architecture of each of the two encoder networks is shown in Fig. 2(b), which includes five fully-connected layers. The dimension of the input and output are, respectively, the number of microstates *n* and the number of macrostate *N*. For example, in the alanine dipeptide dataset, we aimed to lump 100 microstates into 4 macrostates, and therefore there are 100 nodes on the first layer of the encoder and 4 nodes on the last layer of the encoder. In our design, we try to keep the ratio of the number of the nodes between each layer to be similar (see Sec. II.E.5 for the number of nodes for each example). Rectified linear unit (ReLU) ^51,52^ is used as the activation function between each layer, and SoftMax is used for the output so that the output vector resembles a membership function to the macrostates that sums to one. Dropout and max-pooling were used in the first and second layers to avoid overfitting.

The lumping matrix encoded in the neural network after the training procedure could be retrieved by feeding in all the *n* possible one-hot vector assignments (representing *n* microstates) sequentially into the encoder and stacking all the outputs (1 × *N* vectors) together. The obtained *n* × *N* matrix is the fuzzy lumping matrix **A**.

Transpose symmetrization is used when computing the TCMs, for Y-loss computation to ensure detailed balance condition is fulfilled. And the eigenvectors *v*_*T*_ are computed through a singular value decomposition procedure and the update is done using the PyTorch functionality.^53,54^ Scaling and normalization have been done in the singular value decomposition (SVD) step to ensure the eigenvectors of original microstates and reverse projected ones are of the same scale. More details could be found in Appendix B.

### E. System setup and simulation details

#### 1. 1D potential

To illustrate the reverse projection idea using a simple system, we have performed an MD simulation on the 1-D potential presented in Ref^6^:

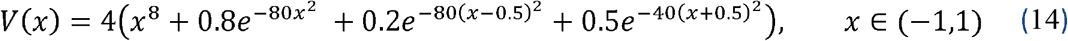

which contains 4 minima in the stated region. Instead of the kinetic Monte Carlo simulation used in the original work^6^, we have performed an NVT MD simulation in the current example. A reflective boundary condition was set on the boundary to prevent the particle from diffusing away from the region. A velocity-Verlet integrator^55^ coupled with Andersen thermostat^56^ (*T=1*, collision frequency = 50 per time unit) was adopted for the NVT simulation. The integration time step is *0*.*0001* time unit and the trajectories is saved every *100* integration steps. So the 10^8^-step simulation resulted in 10^6^ saved frames in total. In this work, we have split the 1D region into 100 equally spaced grids as microstates. The reverse projected eigenvectors are computed based on the two lumping at a lagtime of 50 saving intervals.

#### 2. Alanine Dipeptide

The simulation trajectories for alanine dipeptide are the same as those used in Ref^40^, which is consisted of one hundred 10-ns MD simulations. MD conformations are saved every 0.1 ps and thus there are 100,001 frames per trajectory. In this work, the 10,000,100 MD conformations are split into 100 microstates using k-centers clustering algorithm based on the root-mean-squared distances aligned on all heavy atoms. The microstate Markovian lagtime for this 100-state model is estimated to be 5 ps.

The “PCCA+ lumping” is obtained from the crisp assignment of the PCCA+ fuzzy lumping^38^ result in PyEMMA.^33^ The hierarchical clustering with Ward linkage^57^ is done based on the distance matrix 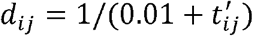, where *t*_*ij*_′ is the element of the symmetrized TCM. MPP lumping^41^ is done with qMin = 0.7 so that 4 states are obtained.

#### 3. 2D potential

In order to examine in detail the behavior of kinetic lumping when the lag time is relatively long and the separation of timescale is relatively small, we have set up an easy-to-visualize single-particle molecular dynamics simulations on the same 2D potential as in Ref ^40^ with the following potential form:

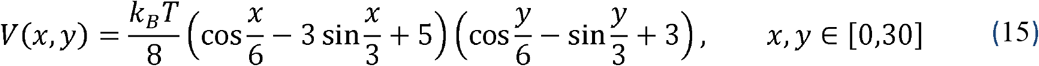

which contains 4 energy minima, and reflective boundary condition set on the boundary to prevent the particle from diffusing out of the region: *x,y* ∈ [0,30]. A velocity-Verlet integrator^55^ coupled with Andersen thermostat^56^ (T=1, collision frequency = 5 per time unit) was adopted for the NVT simulation. The integration time step is 0.001 time unit, and the trajectories is saved every 100 steps. As a result, our 10^9^-step MD simulation trajectories is consisted of 10^7^ saved frames. We have further split the 2D region (*x,y* ∈ [0,30]) into 31 x 31 equally spaced grids, resulting in 961 microstates. The microstate Markovian lagtime of this microstate partitioning is estimated to be 80 saving intervals (8000 integration steps). The PCCA+ lumping is obtained from the crisp assignment of the PCCA+ fuzzy lumping^38^ result in PyEMMA.^33^

#### 4. RNAP loading gate dynamics

The conformational changes of opening and closing of the Clamp conformation of a holoenzyme with the loading of a promoter RNA has been simulated in Ref ^58^. The system contains 543,237 atoms, and the simulation dataset is consisted of 306 200-ns MD trajectories. The all-atom MD conformations were projected into three tICs obtained from the tICA. Then a 100-state microstate model was constructed via k-centers clustering in the three-dimensional tICA space with the tICA lag time of 10 ns. Please refer to Ref ^58^ for more details of the microstate MSM construction and validation.

For this system, the PCCA+ lumping is obtained from the crisp assignment of the PCCA+ fuzzy lumping^38^ via the PyEMMA^33^ software. When performing the kinetic lumping, we have adopted the same microstate assignments as Ref ^58^ but chose a slightly different lagtime of 90ns rather than the lag time of 60ns adopted in Ref^58^. The hierarchical clustering with Ward linkage is done based on the distance matrix 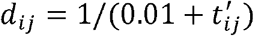, where *t*_*ij*_ ′ is the element of the symmetrized TCM. MPP lumping^41^ is done with qMin = 0.97 so that 4 states are obtained.

#### 5. Parameters of the Neural Networks

As mentioned in Sec. II.D, we aim to keep the ratio of number of nodes between the n-th and the (n+1)-th layer roughly constant, thus we have set up the networks with number of nodes as the following: (1) alanine dipeptide: 100-50-25-16-8-4; (2) 2D potential: 961-300-100-32-12-4; and (3) RNAP loading gate dynamics: 100-50-25-16-8-4.

We made use of the ADAM^59^ optimizer for training, with the learning rate of the network set to 0.001 and a decay rate of 0.99 in each epoch. Dropout probability for the dropout later is chosen to be 0.2, and 30 training epochs were used for all datasets. The batch size of 6000 is used in alanine dipeptide and 2D potential, and batch size of 2000 is used to study the RNAP loading gate dynamics. These batch sizes are chosen by preliminary scanning through the batch sizes ranging from 1000 to 10000, and the tests showed that if the batch sizes are too small or too large, the network might be much harder to converge.

#### 6. Assessing the quality of the lumped models

To compute the GMRQ and metastability of the lumping results, we first randomly divide the dataset into two equal portions as training and test dataset for 30 different times. The lumping matrix is then computed based only on the training dataset and the GMRQ^50^ and metastability are computed for both the training and testing dataset separately for each of the 30 random partitions.

## III. RESULTS AND DISCUSSIONS

We demonstrate the performance of RPnet in three systems: the conformational dynamics of alanine dipeptide in water, a single-particle diffusion on a 2D potential, and the conformational dynamics of the DNA loading gate of a bacterial RNA polymerase.

### A. Alanine Dipeptide

We first demonstrate the performance of RPnet using the commonly used benchmarking example: the alanine dipeptide in water. The sampled conformations are grouped into 100 microstates (see Methods for details). As there exists a stable gap between the 3^rd^ and 4^th^ slowest implied timescale (Fig. 3(h)), we lumped the 100 microstates into 4 macrostates, which is also consistent with previous studies^40,60^. As shown in Fig. 3 (a & c), RPnet correctly identifies the four metastable states 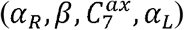. The implied timescales of the resulted lumping (Fig. 3(g)) also reproduce those of the microstates reasonably well. We can also clearly visualize the optimization process of RPnet in Fig. 3(f). Although the initial Y-loss is high (at 0.18755), after around 20 epochs, the Y-loss has gone down to 0.00127, which is the same as the Y-loss calculated for PCCA+ and MPP (see Fig. S1a for the corresponding Y matrices), consistent with same lumping results shown in Fig. 3 (b,c,e). In this example, RPnet displays comparable performance with other popular kinetic lumping methods (e.g., PCCA+, Hierarchical clustering using Ward linkage, and MPP). Specifically, all of these methods can correctly identify 4 macrostates containing largely similar microstate assignments (Fig. 3(b-e)). Because alanine dipeptide is a well-studied system containing clear separation of timescales, it is expected that all these kinetic lumping methods yield comparable results. In the next example, we will apply RPnet on a more challenging system with less clear separation of timescales, where we show that RPnet can robustly identify the metastable states and outperform other kinetic lumping methods.

**FIG. 3.**
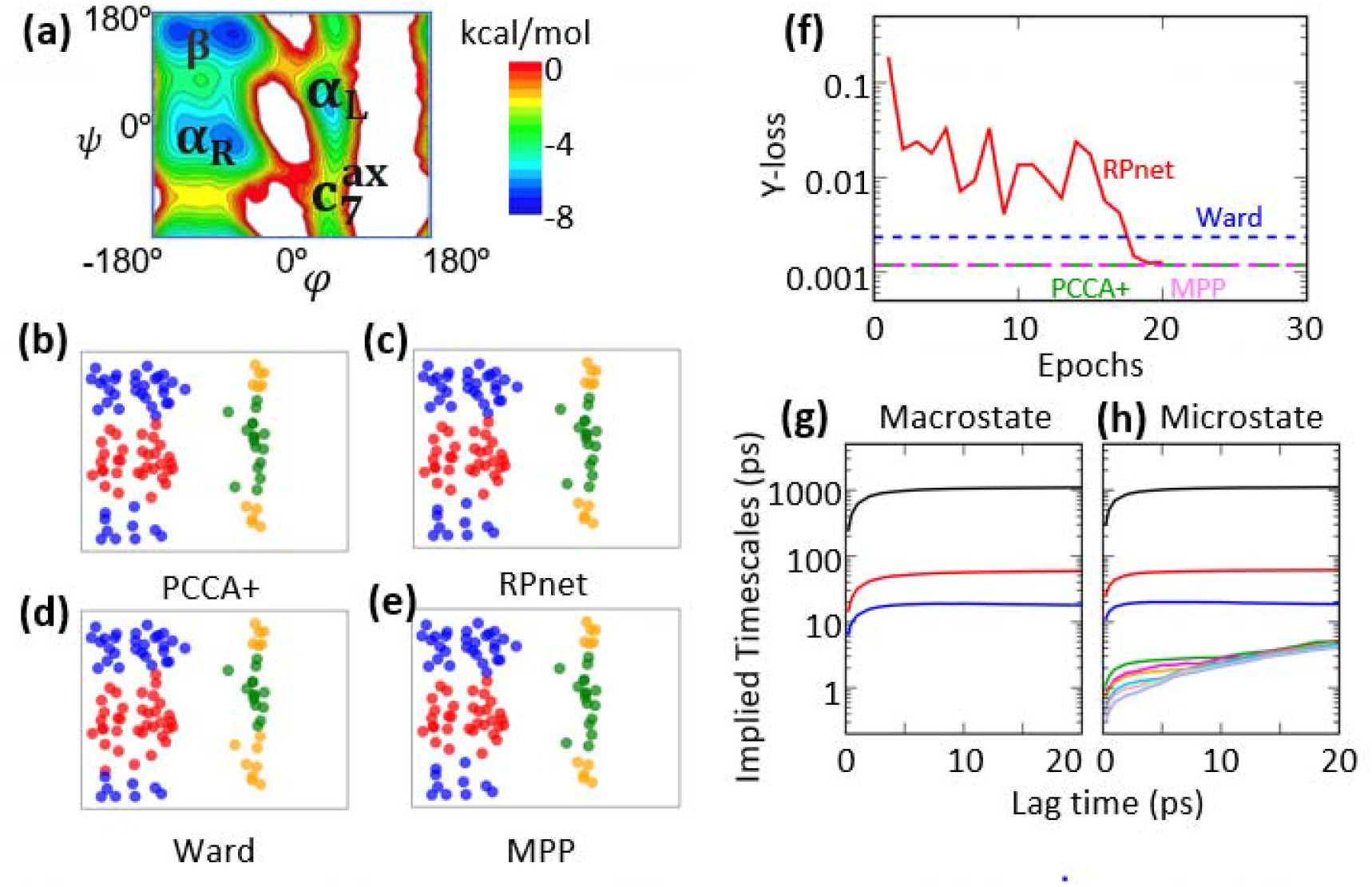
Performance of RPnet on the alanine dipeptide system. (a) The Ramachandran plot of the alanine dipeptide. (b-e) Lumping results by 4 different algorithms. (b) PCCA+. (c) RPnet. (d) Hierarchical clustering with Ward linkage. (e) MPP. (f) Evolution of the Y-loss of RPnet over 30 epochs, with referenced to the Y-loss of the other three lumping. (g-h) Implied timescales of the (g) 4 macrostates resulted from RPnet, and (h) original 100 microstates.

### B. 2D potential

Although the alanine dipeptide provides as a good benchmark system for the development of kinetic lumping algorithms, it is often too simple to differentiate the performance of difference algorithms. In particular, the separation of timescale of the constructed MSM might be less clear in more complex systems, and this may impose challenges to eigenspectrum-based lumping methods such as PCCA^35^ or its variants^36–38^ due to the numerical instability. To investigate this situation, we have designed a 2D potential, where its 4 minima can be analytically identified as the gold standard of lumping results (see Eq. (15)). We have divided the XY-space as shown in Fig 4 into 31 × 31 equally spaced grids, thus forming a 961-state microstate model. We then tried to lump the 961-microstate model into a 4-macrostate model (see Methods for more details). To create a situation with less clear separations of timescales, we have adopted a relatively long lagtime (3000 saving intervals) to construct the macrostate model when analyzing the single-particle diffusion on this 2D potential.

**FIG. 4.**
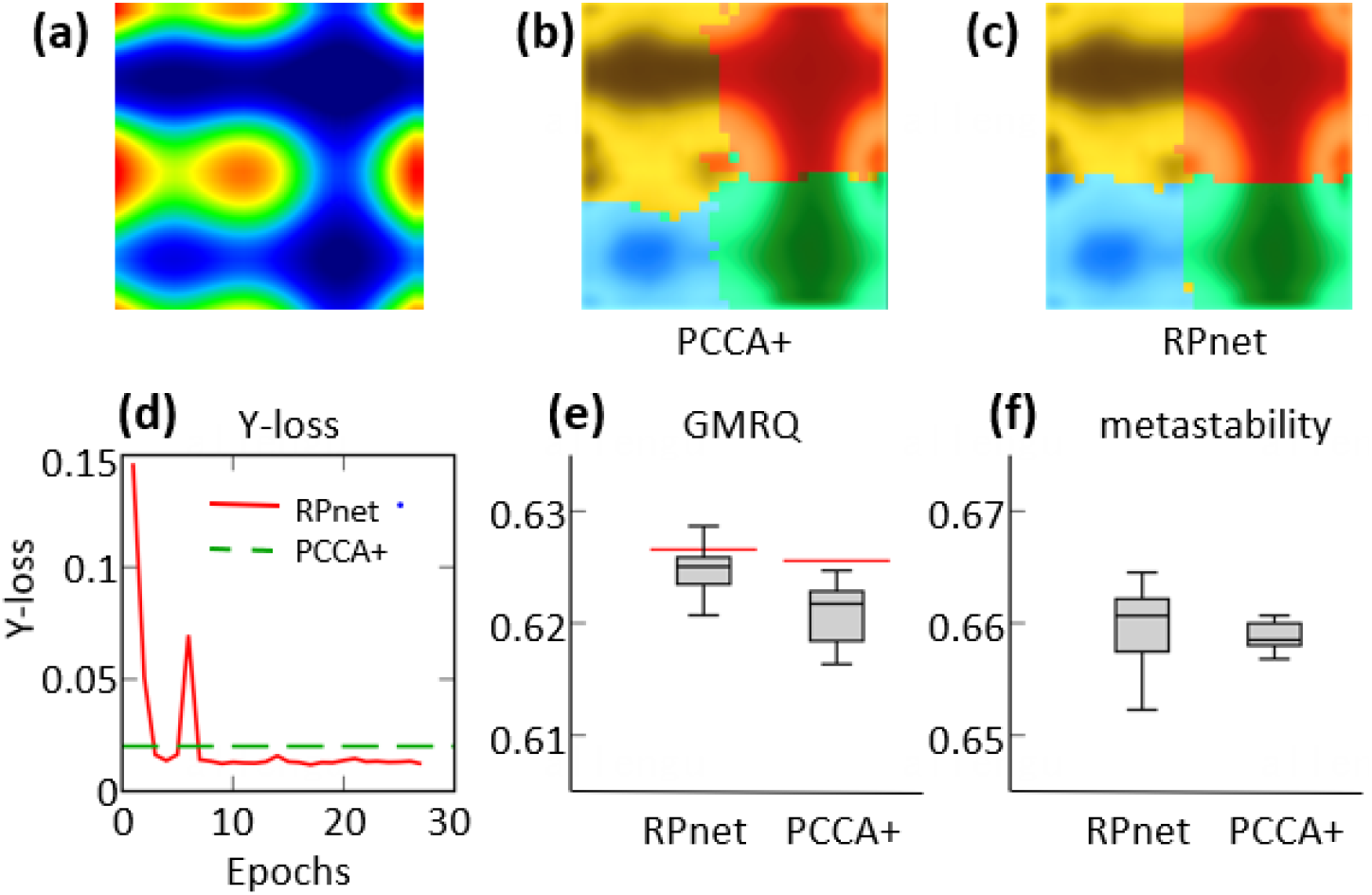
Performance of RPnet on the 2D potential system. (a) The potential energy surface used in this test. (b-c) Lumping results at lagtime *τ* = 3000 for (b) PCCA+, and (c) RPnet. (d) Evolution of the Y-loss of RPnet over 30 epochs, with referenced to the Y-loss of the PCCA+ lumping. (e-f) Comparison of (e) Generalized Matrix Rayleigh Quotient (GMRQ) and (f) metastability of the two lumping methods, redline denotes the training mean and the box-and-whisker plot denotes the distribution of 30 test sets (see Sec. II.E.6 for details).

As shown in Fig. 4(b), the state boundary of the PCCA+ becomes fuzzy at this lagtime (*τ* =3000) and some microstates belonging to the free energy minima in blue) were mis-assigned to the yellow macrostate, while our RPnet correctly separate the blue state from the yellow state (see Fig. 4c). The optimization process of our RPnet is presented in Fig. 4(d), which clearly indicates that the RPnet achieved a lower Y-loss (∼0.01224) than that of PCCA+ (see Fig. S1b for the corresponding Y matrices and Fig. S2 for the implied timescales). In addition to the Y-loss value, we have also evaluated the quality of lumped macrostate models using two other criterions: the metastability and the GMRQ. Metastability measures the average probability of self-transitions between macrostates, and a high metastability usually indicates a good separation of slow inter-state dynamics and fast intra-state dynamics. GMRQ assesses the quality of the macrostate models via a cross-validation process with an objective function based on the variational principle of conformational dynamics. As shown in Fig. 4(e), the lumped model from RPnet results in higher GMRQ and metastability, indicating that the macrostate model obtained from RPnet is better than that of PCCA+. These observations suggest that PCCA+ could suffer from the instability of eigenvector components and thus generate fuzzy state boundaries and leads to lowered GMRQ and metastability. We anticipate that this situation may be more prominent in complex biological systems, while our RPnet can yield more robust results in kinetic lumping.

### C. Dynamics of the clamp domain of a bacterial RNA polymerase

After applying RPnet to two simple systems, we here proceed to a realistic biological system: i.e. the conformational conformation of bacterial RNAP clamp opening/closing motion that we reported recently^58^ (see Methods for details). As shown in Fig. 5a, the four macrostates of this system correspond to the open state (State S1, yellow dots), the closed state (State S4, green dots), as well as two partially closed states differing by the switch 2 region; partially closed with α-helix switch 2 (State S2, blue dots) and *π* -helix switch 2 (State S3, red dots). The switch 2 region is a helical structure under the clamp domain which is crucial for the clamp movement^58^. In the PCCA+ based four-state model reported before, the transitions between S1/S2 or S3/S4 only takes 10∼100 μs, while the transitions between the two partially closed state S2/S3 would take 2 ms, indicating a huge gap between S2 and S3. We use the tICA projection (tICA lag time = 10 ns) for the visualization of different macrostate lumping.

**FIG. 5.**
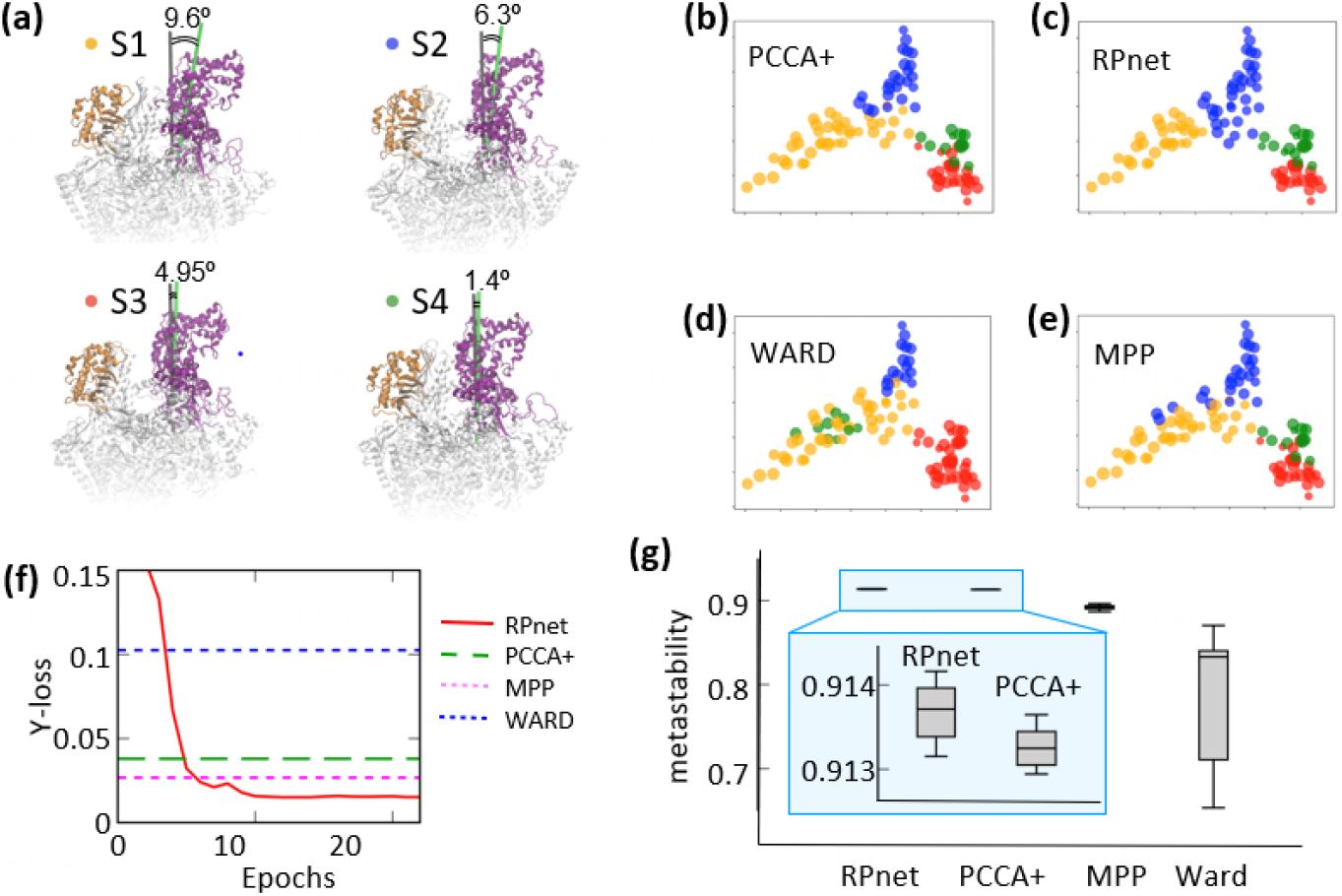
Performance of RPnet on the RNAP clamp motion. (a) representative conformations of the four metastable states shown as yellow (state S1), blue (state S2), red (state S3) and green (state S4) dots of (b-e). The Clamp open/close angles are also presented in the snapshots of metastable states. (b-e) Lumping results at lagtime *τ* = 90 ns for (b) PCCA+, (c) RPnet, (d) Hierarchical clustering with Ward linkage, and (e) MPP. (f) Evolution of the Y-loss of RPnet over 30 epochs, with referenced to the Y-loss of the other three lumpings. (g) Comparison of metastability of all four lumping methods. The box-and-whisker plot denotes the distribution of 30 test sets (see Sec. II.E.6 for details).

As shown in Fig. 5(c), the four macrostates obtained by RPnet can be well separated in the projection of microstate centers onto the top two tICs, indicating that the macrostate boundaries clearly reflect the slowest dynamic modes of the system. Furthermore, the Y-loss score of the RPnet is the lowest (see Fig. 5(f)), and it also has the highest metastability among all the methods (see Fig. 5(g)). This state assignment is fully consistent with the previous study^58^, which was produced by PCCA+ with a shorter lag time (see Methods for details).

Close examination of the Y-loss of the four assignments would reveal that Y-loss does correlate well with the quality of different lumpings (see Fig. S1c for the corresponding Y matrices). For example, the macrostate in green color is totally mis-assigned in the model obtained from the hierarchical clustering with Ward linkage (see Fig 5(d)), and this model also yields the highest Y-loss value. For PCCA+ (see Fig. 5(b)) and MPP (see Fig. 5(e)), both methods generate models in overall good quality, however, a few microstates at the state boundaries are not correctly assigned especially between the green and red macrostate. As a result, PCCA+ and MPP substantially improve in the Y-loss scores compared to the Ward linkage clustering, but they still produce models with higher Y-loss score than our RPnet.

Taking all the results together, we show that RPnet can serve as a powerful kinetic lumping algorithm. The novelty of RPnet lies in its objective function (or loss function), which is based on the reverse projection of conformational dynamics from a macrostate model to a microstate model. In particular, the Y-loss function directly examines the similarity between the transition modes before and after kinetic lumping. This is distinct from the variational principle based objective function, which is focus on optimizing the slowest timescales (i.e., eigenvalues of the TPMs). We show that at situations in which the separation of timescales between intra- and inter-state transitions is not clear, our Y-loss function could robustly identify lumpings with correct and cleaner state boundaries. In contrast, algorithms that are based on eigen-decompositions of TPMs (e.g. PCCA+) may become more sensitive to numerical instabilities under those conditions (e.g. in the 2D-potential example in Fig. 4, see also Fig S3 for the analysis on robustness).

Our design of the loss function is inspired by a previously developed projection operator framework by Hummer and Szabo^48^. In that study, they make use of the projection operator framework to construct a transition matrix that can best describe the dynamics of a prespecified macrostate lumping, by matching the time-dependent occupation-number correlation function of microstates and macrostates. Although this method would be able to extrapolate the population evolution (in terms of time-dependent occupation-number correlation functions) of multistate kinetics reasonably well, we anticipate that it may not perform well when applied to distinguish good and poor lumpings as all the kinetic lumpings would result in reasonably good approximation of the evolution using their method. Our work gains inspiration from the projection operator framework developed by Hummer and Szabo^48^, but our aim is not at the extrapolation of kinetics, and instead we further designed a reverse projection to evaluate the quality of lumping at the same time. As a result, our RPnet is not simply a tool to predict long timescale kinetics, but also produce the optimized state partitioning.

We note that although the Y-loss in our RPnet scheme itself can already be used to judge the quality of lumping, the use of encoder network provides an efficient approach to search for lumped models. Some early attempts of kinetic lumping^11,61^ optimizes lumping assignments through repeated trials and Monte-Carlo type optimization, in which the improvement between iterations are often relatively small and the optimization is easily trapped into local minima. With the use of a neural network, not only the combination of SoftMax activation functions could approximate the complex loss function landscape, it also allows a parallelized nonlinear search via backpropagation and quickly identify a good solution.^62^ In fact, the neural network also encodes the key representation of the complex system dynamics, which not only facilitates the backpropagation update,^63^ but also allows easy extension of the neural network architecture for other purposes like fuzzy lumpings.

## IV. CONCLUSION

In this work, we have developed a kinetic lumping algorithm: RPnet, that combines a physics-based loss function and the optimization using neural network. Inspired by the projection operator framework in statistical mechanics, we have designed a reverse projection scheme for comparing the transition modes among original microstates and lumped macrostates, which allows the quantification of lumping quality that is truly based on the underlying physical process. Using our proposed Y-loss based on the reverse projection scheme, we have also set up a neural network that allows an automatic optimization of lumping. Combining the two parts, we have obtained our RPnet framework, that allows an automatic method that can give rise to physically sound lumping. As demonstrated by the three systems in the text, this framework can have good performance across a wide range of systems. The projection-operation-based loss function also provides an alternative line of thought for the evaluation of macrostate lumping quality and should give new insight to the identification of slow dynamics in complex systems. We anticipate that our method holds promise to be widely applied in the MSM construction to study protein functional dynamics.^28^

The source code of RPnet is available for download at: https://github.com/ghl1995/BpNet-lumping.

## SUPPLEMENTARY MATERIAL

See supplementary material for the details of Y-matrices in different cases and implied timescales of the 2D potential example.

## ACKNOWLEDGMENTS

We thank Dr. Gerhard Hummer for the inspiration and fruitful discussions. X.H acknowledges the support from the Padma Harilela Endowment Fund.

## Appendix A Theory of Reverse Projection

Prove: idempotency ℙ^2^ = ℙ, ℚ^2^ = ℚ, ℙℚ = 0, and also the identity **A**^T^ ℙ ≡ **A**^T^

A well-known property of the projection operators ℙ and ℚ is ℙ^2^ = ℙ, ℚ^2^ = ℚ, ℙℚ = 0. This could be proved by 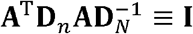. From Eq. (3) we have *P*_*eq*_ = **A**^T^*p*_*eq*_, which can be written as the matrix form: **D**_*N*_ = **A**^T^**D**_*n*_**A**, therefore 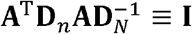. Then we will have,

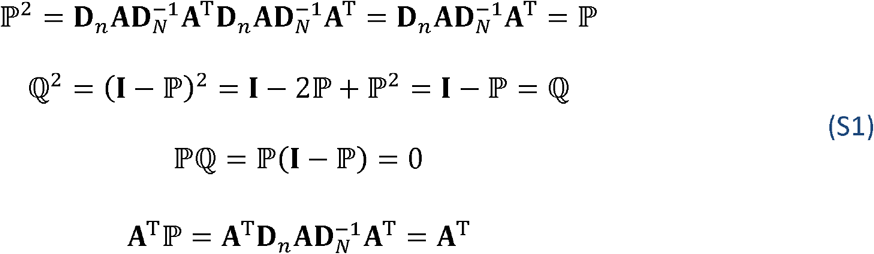

which proves the proposition.

A simple understanding of Eq. (11) can be made with the variational principle, *λ*_*M*_ ≤ *λ*_*T*_. In the spectral space that used for variational principles, the eigen decomposition of **T**(t) is written as: **T*v***(*i*) = ***v***(*i*)*λ*_*T*_ and ***ϕ***(*i*)^T^**T**= *λ*_*T*_***ϕ***(*i*)^T^, where 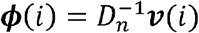. At the same time, the eigen decomposition of **M**(*t*) is written as: **M*V***(*i*) = ***V***(*i*)*λ*_*M*_ and **Φ**(*i*)^T^**M**=*λ*_*M*_ **Φ** (*i*)^T^, where 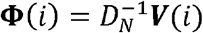. Simply, when the top eigenvectors satisfy 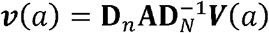 and ***ϕ***^T^(*a*) = **Φ**^T^(*a*)**A**^T^, we will achieve the variational principle:

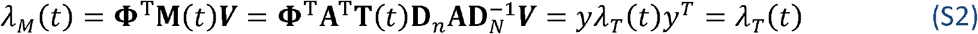

**T**(*t*) ≡ ***v****λ*_*T*_(*t*) ***ϕ***^T^. *y* is the overlapping matrix defined in Eq. (12).

A rigorous proof is shown as follows. First, straightforwardly from Eq. (5),

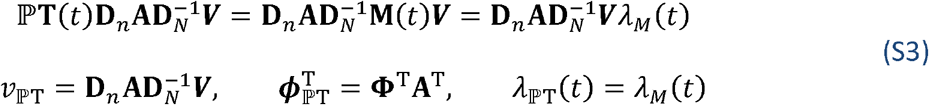

Indicating the right eigenvector and eigenvalue of ℙ**T**(*t*) are 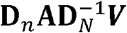 and *λ*_*M*_(*t*), respectively. Secondly, when both microstate and macrostate model have master equations,

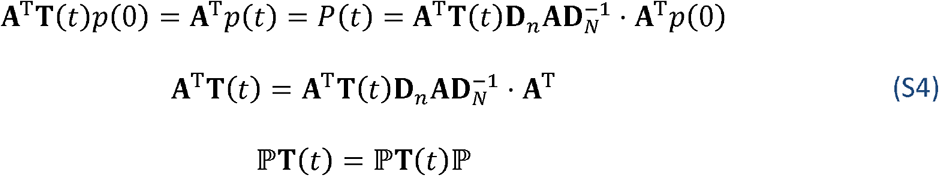

Then make use of the relation ℙ**T**(*t*) = ℙ**T**(t) ℙ,

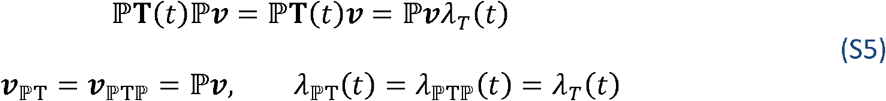

Indicating that the right eigenvector and eigenvalues of ℙ**T**(t)ℙ or ℙ**T**(t) should be ℙ***v*** and λT(t) respectively. Now combine the conclusion Eq. (S3) and (S5), we have:

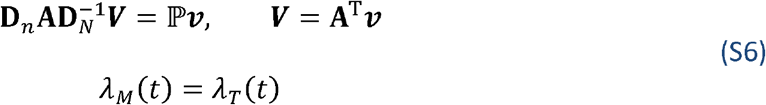

Then we will prove ***v*** = ***v***_ℙT_:

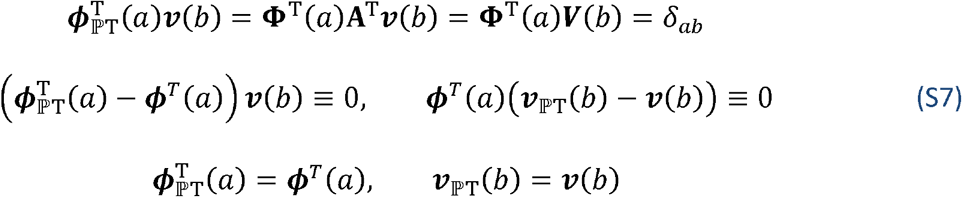

Where the above equations only apply for top eigenmodes: 0 < *a, b* ≤ *N*. Therefore,

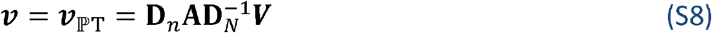

Which is Eq. (11).

## Appendix B RPnet algorithm

The following Algorithm 1 shows how we implement the RPnet in the network. In neural network, back propagation cannot support eigen decomposition, thus, we need to use SVD to replace it. The key idea is to borrow the relation of singular vectors of transition count matrix and eigenvector of transition probability matrix. Specifically, according to Eq. (S3) the relation between singular vector ***P*** and eigenvector ***ϕ*** is: ***ϕ*** = **D**^−0.5^***P***:

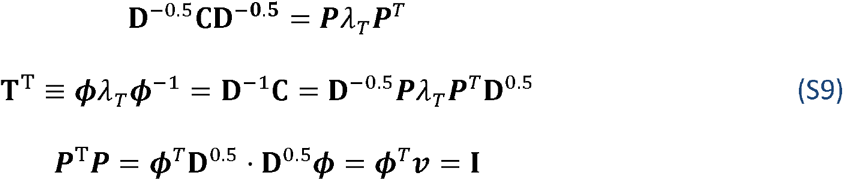

### Algorithm 1 RPnet

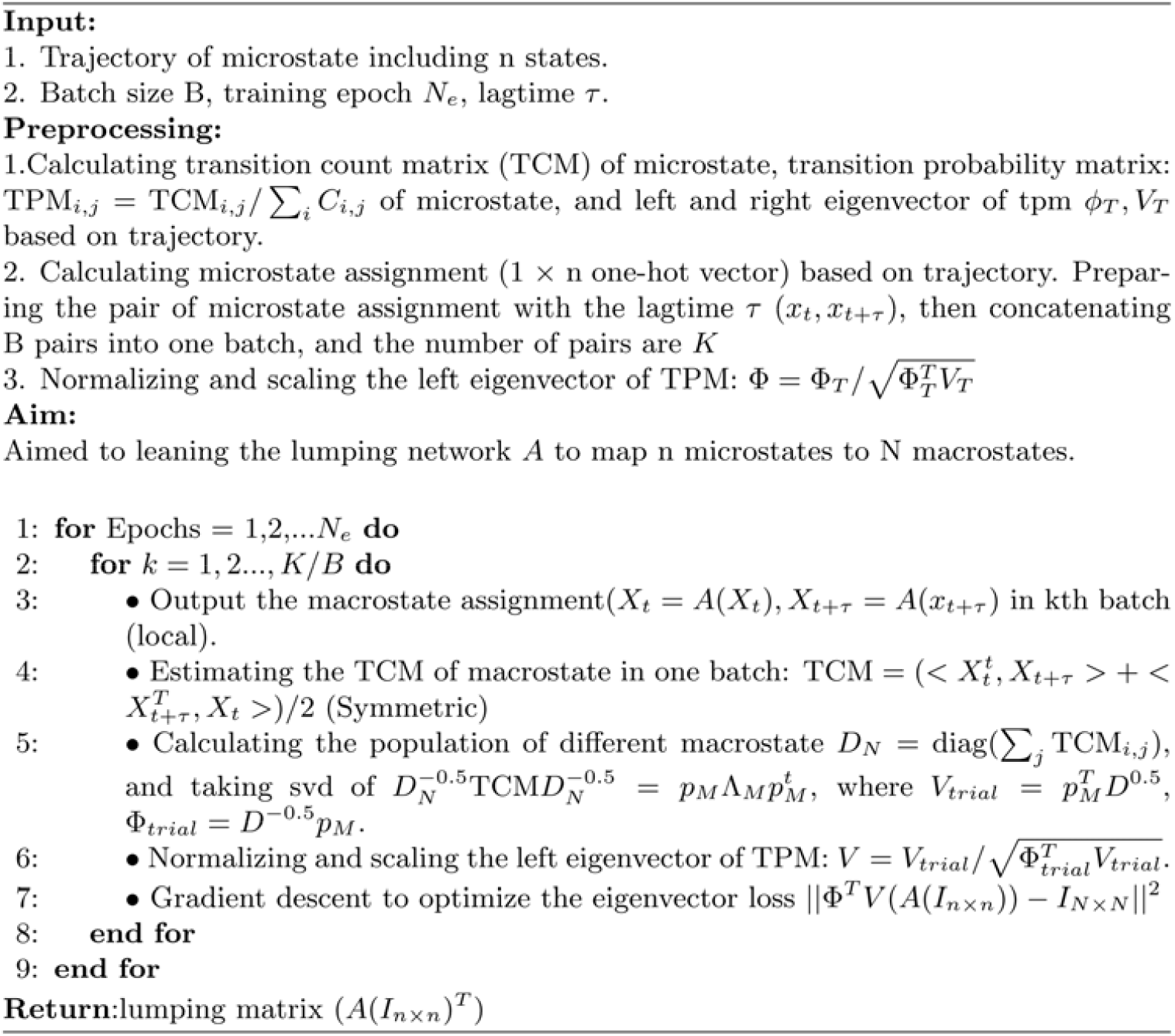

