## Supplementation materials for "RPnet: A Reverse Projection Based Neural Network for Coarse-graining Metastable Conformational States for Protein Dynamics"

‡ Hanlin Gu, Wei Wang and Siqin Cao contribute equally to this work.

### Supplementary Text

#### 1. Comparison of Y matrix obtained from different kinetic lumping methods

In this section, we present the underlying Y matrices that are used to compute the Y loss of the corresponding cases, namely the *Alanine Dipeptide* (lagtime of 5ps), 2D potential (lagtime of 3000 saving intervals), and RNAP (lagtime of 90 ns) respectively (Fig. S1a-S1c). We used the deep blue colour to represent $y_{ij} = 1$ and light blue to represent $y_{ij} = 0$.

For the *Alanine Dipeptide* case (Figure S1), all methods result in a Y matrix with close resemblance to the identity matrix, with the diagonal elements all close to 1 and off-diagonal elements all close to 0. In fact, RPnet, PCCA+ and MPP give exact same lumping results, and so their Y matrices are also the same.

For the 2D potential system (Figure S2), the Y matrix corresponding to RPnet has larger diagonal elements and smaller off-diagonal elements when compared to that resulted from the PCCA+, consistent with the better state boundary partitioning of RPnet.

For the case of RNAP (Figure S3), the Y matrix of RPnet is again closest to identity matrix. It can also be seen that the Y matrix corresponding to hierarchical clustering with Ward linkage actually has a mixing between the third and fourth eigenvectors, which correspond to the erroneous state partitioning shown in FIG. 5.

#### 2. Comparison of the implied timescales of macrostate-MSMs generated by different kinetic lumping methods

Figure S4 shows the implied timescale of the 2D potential dataset. Panel (a) presents the 9 slowest implied timescales of the microstate model. The implied timescales of the lumped macrostate models from RPnet and PCCA+ are shown in panel (b) and (c), respectively. As shown in Figure S4, the implied timescales obtained from 4-macrostate MSMs generated by RPnet and PCCA+ are both consistent with the three slowest implied timescales predicted by the microstate-MSM.

#### 3. Stability of RPnet in different lagtime

We have presented in the main text that our RPnet method performs better than other methods in several specific lag times. We will hereby show that our RPnet approach is also robust, where Y-loss is always low and stable. Figure S5 displays the Y-loss values computed at different lag times in the three systems. In order to demonstrate the stability in the performance of RPnet, we compare our method to the PCCA+. The result demonstrates that RPnet Y-loss is significantly less sensitive to the value of the lag time compared to PCCA+. Furthermore, we show that the Y-loss values of RPnet are always lower than those from PCCA+, even though PCCA+ can achieve comparable performance with RPnet in some specific lag times (see Figure S5).

### Supplementary Figures:

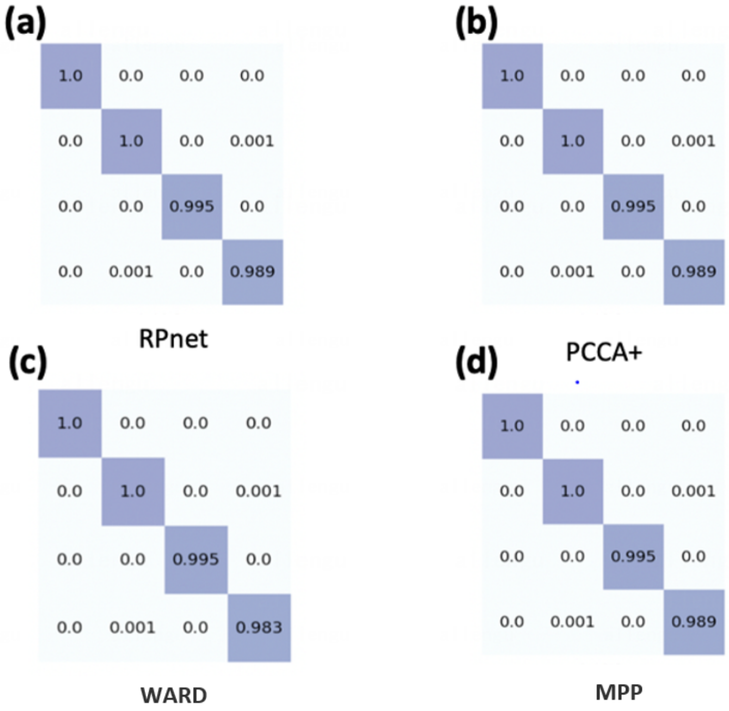


**Figure S1:** The Y matrix of *Alanine Dipeptide* built with the lagtime of 5ps. The microstate model has 100 states, and the macrostate models have 4 states. In (a-d), the macrostate models are generated by RPnet, PCCA+, hierarchical clustering with Ward linkage, and MPP, respectively.

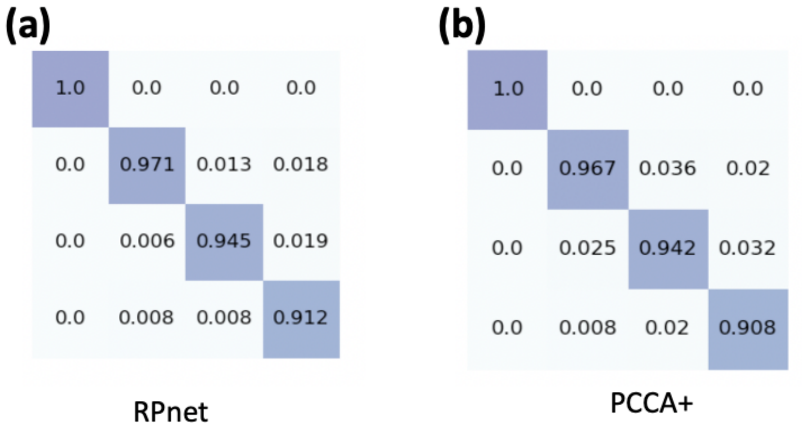


**Figure S2:** The Y matrix of 2D-potential with the lag time of 3000 steps. The microstate model has 961 states while the macrostate models have 4 states. The macrostate models are generated by: (a) RPnet and (b) PCCA+, respectively.

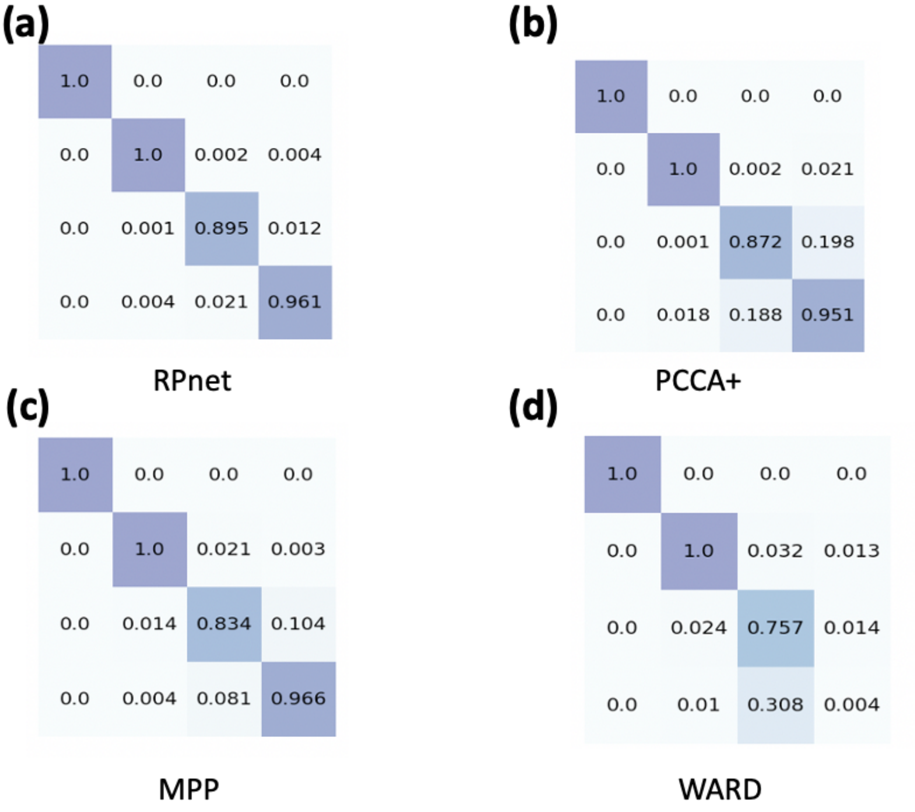


**Figure S3:** The Y matrix of RNAP with 90 ns lag time. The macrostate model has 100 states, while the macrostate models have 4 states. In (a-d), the macrostate models are generated by RPnet, PCCA+, MPP and hierarchical clustering with Ward linkage, respectively.

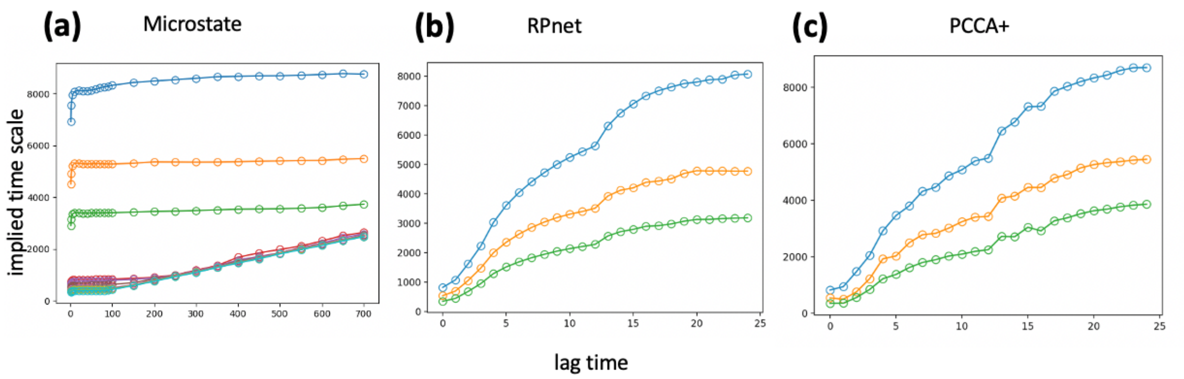


**Figure S4:** The implied time scales of 2D potential system with different lag time. (a) is the implied time scale of Microstates. (b) is the implied time scale of macrostates generated by RPnet, (c) is the implied time scale of macrostates generated by PCCA+.

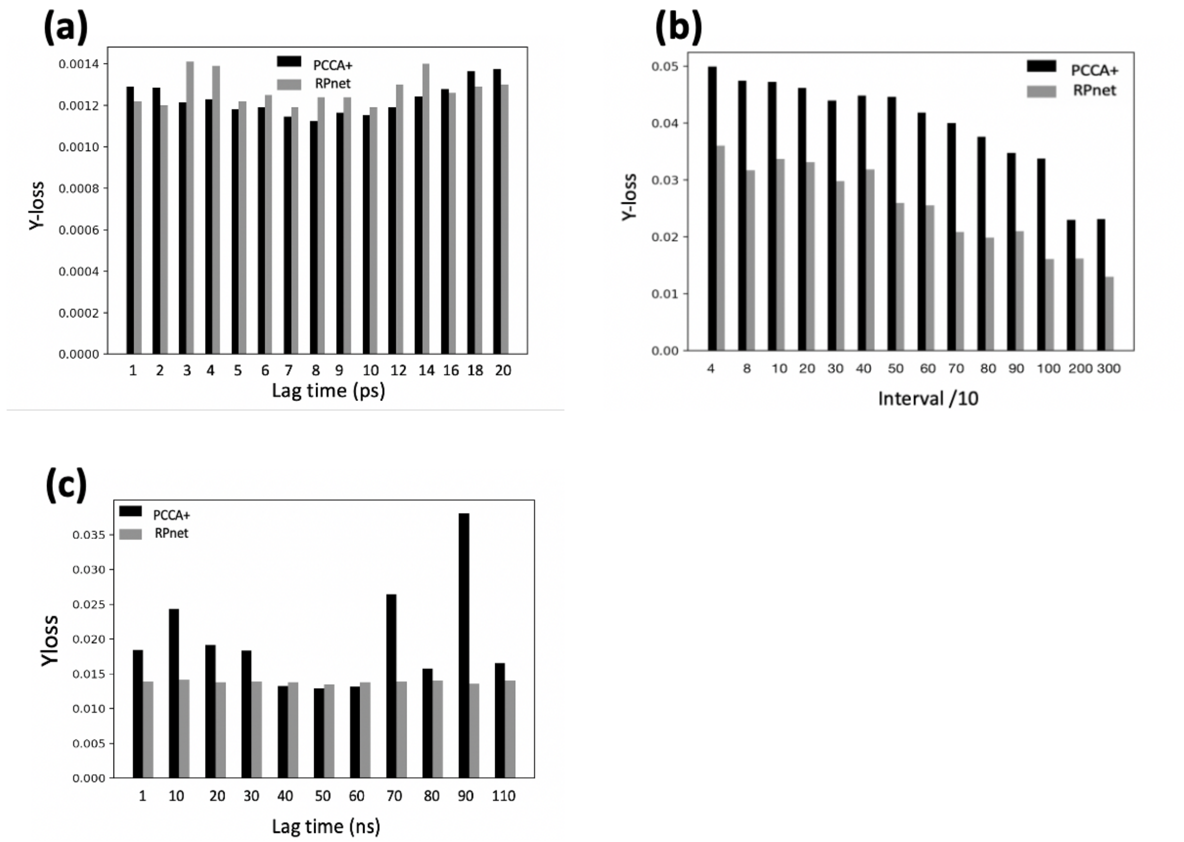


**Figure S5:** The Y-loss result with different lag time. (a) is the Y-loss change in the *Alanine Dipeptide*. (b) is the Y-loss change in the 2D potential. (c) is the Y-loss change in the RNAP.
